# When good mutations go bad: how population size can change the direction of natural selection

**DOI:** 10.1101/2021.04.19.440541

**Authors:** Yevgeniy Raynes, Christina L. Burch, Daniel M. Weinreich

## Abstract

Classical evolutionary theory holds that the efficiency, but not the direction, of natural selection depends on population size. In small populations, drift overwhelms selection, rendering all fitness-affecting mutations selectively neutral. Yet, beneficial mutations never become deleterious and deleterious mutations never become beneficial. Remarkably, several mutations, including in modifiers of recombination and mutation rate, have now been shown to be favored at some population sizes but disfavored at others, challenging established theory. Previously, we have designated this phenomenon sign inversion. Here we show that, unlike selected mutations in the classical framework, mutations susceptible to sign inversion confer both fitness costs and fitness benefits, that vary among their carriers. Furthermore, all such mutations can be classified based on whether their effects differ between or within mutant lineages. Using computer simulations, we demonstrate that both between-lineage and within-lineage variability can cause sign inversion and elucidate the common underlying mechanism. Our results confirm that variability in the sign of selective effects is necessary for sign inversion, which occurs because drift overwhelms selection on carriers bearing the cost and carriers enjoying the benefit at different population sizes.

---

The evolutionary fate of a new mutation reflects both the deterministic influence of natural selection and the stochastic influence of genetic drift. Selection applies a biased, asymmetric pressure on mutant frequencies in response to heritable differences in individual lifetime reproductive output, or fitness: upward for beneficial mutations that increase fitness and downward for deleterious mutations that decrease it. Genetic drift exerts an unbiased, diffusive pressure on mutant frequencies, which reduces selection’s efficiency. Accordingly, evolutionary theory often adopts a probabilistic approach, focusing on the probability that a mutation will eventually reach fixation (*P*_fix_) and comparing it with the fixation probability by drift alone (*i.e*., *P*_fix_ of a selectively neutral mutation). The *P*_fix_ of a neutral mutation is equal to 1/*N*, the reciprocal of population size *N* [1]. To control for the population size dependence of the neutral expectation, we will generally focus attention on a mutation’s *N*·*P*_fix_, i.e., *P*_fix_ normalized by the neutral benchmark, 1/*N*. *N*·*P*_fix_ of a neutral mutation is equal to 1 (Fig. 1A).

**Figure 1:**
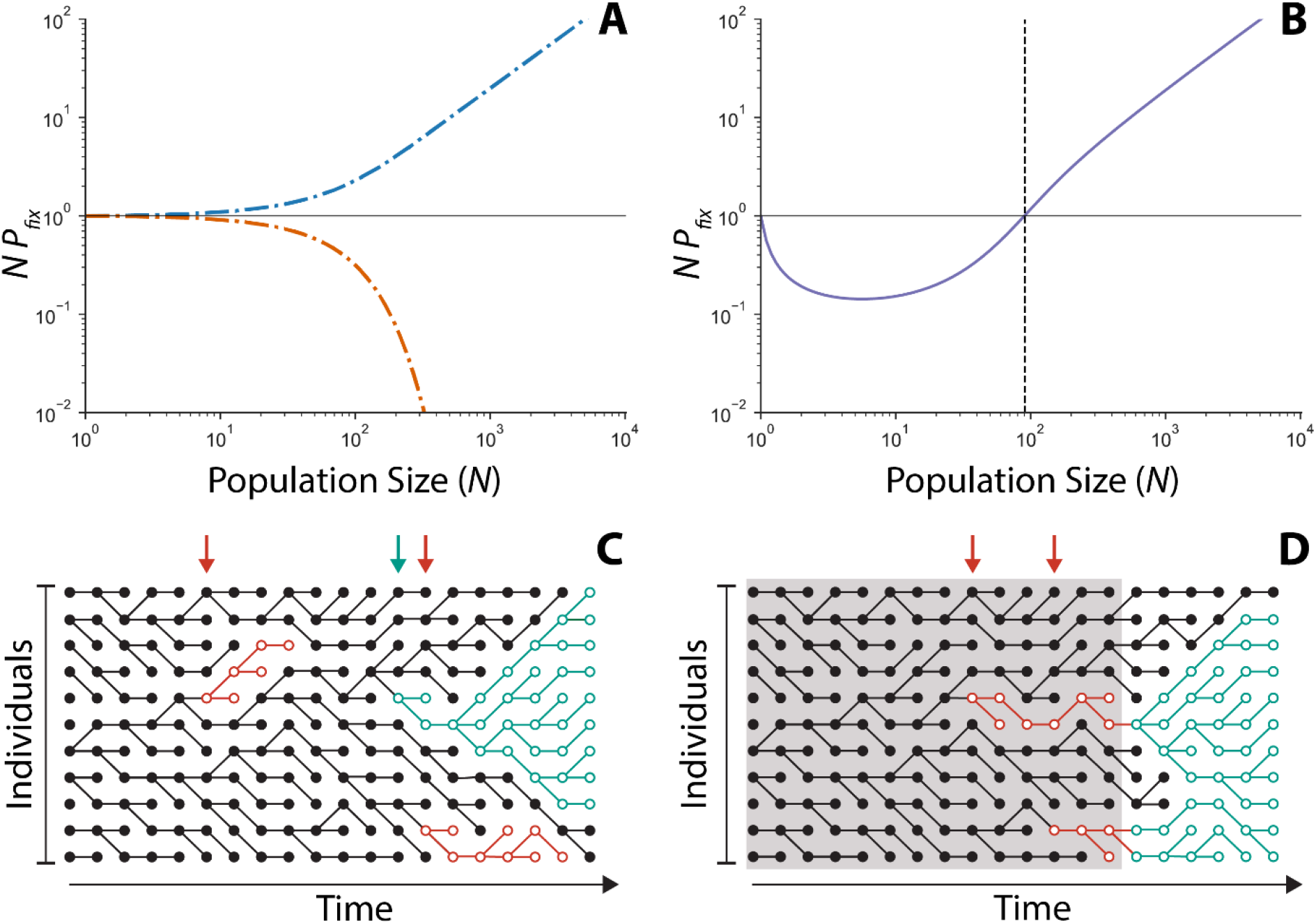
Sign inversion and sign variability. **A)** *N*·*P*_fix_ (given by Eq. 1) of lineage-invariant mutations never crosses 1. *N*·*P*_fix_ of a beneficial mutation (blue, *s* = 0.01) is greater than 1 at all *N*>1. *N*·*P*_fix_ of a deleterious mutation (orange, *s* = −0.01) is less than 1 at all *N*>1. Both equal to 1 at *N*=1. *N*·*P*_fix_ of a neutral mutation (black, *s* = 0) is always 1. **B)** *N*·*P*_fix_ of a sign-variable mutation (purple) exhibits sign inversion when it crosses 1 at *N*crit (indicated by the vertical line). Here, the sign-variable mutation increases both the expectation and the variance in offspring number (see Table 1, computed with Eq. 13 from [5]). The mean ± variance in offspring number for the mutant and the wild type are 1.01 ± 1.0 and 1.0 ± 0.1, respectively. **C** and **D** show genealogies (lines) of haploid individuals (circles) across time (horizontal axis). Carriers of the wild-type allele shown in black. Carriers of sign-variable mutations shown in red and teal. A lineage comprises a complete genealogy of a mutation from its first carrier to fixation or loss. **C)** Between-lineage sign variability manifests between carriers in different lineages. The sign-variable mutation appears 3 times (arrows), giving rise to two lineages bearing the cost (open red circles) and one endowed with the advantage (open teal circles). **D)** Within-lineage variability manifests among carriers (open circles) within a lineage. The sign-variable mutation appears twice (arrows). Here variability is modulated by time: both sign-variable lineages bear an initial cost (red) during the time indicated by the gray background but eventually develop a fitness advantage (teal). As discussed in the main text, within-lineage variability can also be modulated by frequency.

Under classical evolutionary theory, *N*·*P*_fix_ of a beneficial mutation is always greater than or equal to that of a neutral mutation (i.e., *N*·*P*_fix_ ≥ 1), while *N*·*P*_fix_ of a deleterious mutation is always less than or equal to that quantity (i.e., *N*·*P*_fix_ ≤ 1). This is because in the classical framing the fitness effects of all selected mutations are assumed to be lineage-invariant [2] or, in other words, identical in all carriers. Accordingly, the evolutionary fate of any mutation can be computed recursively from its expected per-generation effect on the frequency of its carriers [2]. On this premise, Kimura [1] found the fixation probability of a mutation in a haploid Wright-Fisher population of size *N* to be

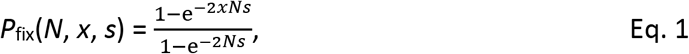

where *x* is the initial frequency of the mutation and its fitness effect relative to the resident wild type is given by the selection coefficient *s*. Kimura’s expression for *P*_fix_ (normalized by 1/*N*), plotted as a function of population size in Fig. 1A, illustrates the classical understanding of the relative influence of selection and drift on the fate of a lineage-invariant mutation. Whereas selection is most efficient in large populations, the influence of drift is magnified in small populations. Consequently, all fitness-affecting mutations – whether beneficial or deleterious – behave progressively more like the neutral benchmark as population size declines. Critical for our purposes, though, the fixation probability of lineage-invariant mutations never crosses the neutral benchmark (Fig. 1A).

We recently discovered that mutations affecting recombination and the mutation rate violate these classical expectations [3, 4] – both have a probability of fixation greater than the neutral benchmark at some *N* but less than that at other *N*. We have designated this phenomenon sign inversion [3], and, subsequently, became aware of two other published examples [5, 6] (Table 1). Algebraically, sign inversion means that the mutation’s *N*·*P*_fix_ crosses 1 (e.g., Fig. 1B) at some intermediate population size, which we have previously designated *N*_crit_ [3]. Necessarily, in all cases *N*·*P*_fix_ converges to 1 in the limit as *N* approaches 1 (compare Fig. 1A and 1B), as in classical theory.

**Table 1:** Published cases of sign inversion involve alleles with sign-variable effects.

|  |  |  |  |  |  |
| --- | --- | --- | --- | --- | --- |
| | $N \cdot P_{\text{fix}}$ | | Sign-variable effects | | |
| Mutation's proximal effect | when $N < N_{\text{crit}}$ | when $N > N_{\text{crit}}$ | Cost | Benefit | Ref. |
| **Between-lineage sign variability** |  |  |  |  |  |
| *Increased expectation and variance in offspring number* | $N \cdot P_{\text{fix}} < 1$ | $N \cdot P_{\text{fix}} > 1$ | Stochastic lineage loss | Increased mean offspring number | [5] |
| *High mutation Rate* | $N \cdot P_{\text{fix}} < 1$ | $N \cdot P_{\text{fix}} > 1$ | Genetic linkage to deleterious mutations | Genetic linkage to beneficial mutations | [3] |
| **Within-lineage sign variability** |  |  |  |  |  |
| *Recombination* | $N \cdot P_{\text{fix}} < 1$ | $N \cdot P_{\text{fix}} > 1$ | Deleterious recombination load | Reduced deleterious mutation rate | [4] |
| *Tit-for-tat cooperation* | $N \cdot P_{\text{fix}} > 1$ | $N \cdot P_{\text{fix}} < 1$ | Decreased fitness v. cheaters | Increased fitness v. other cooperators | [6] |

An examination of the mutations known to exhibit sign inversion reveals that, unlike lineage-invariant mutations, all are characterized by fitness effects that vary among their carriers. Specifically, in each case we find that some carriers of the mutation experience a selective benefit while others are burdened with a selective cost (Table 1). Thus, not only do these mutations exhibit lineage-variable fitness effects (as defined in [2]), the actual sign of their fitness effects varies among individual carriers. Accordingly, we will refer to them as having sign-variable effects. Previously, we found that mutator mutations (which elevate the mutation rate) are susceptible to sign inversion because they have sign-variable effects: individual mutator carriers can indirectly experience either the positive selection acting on linked beneficial mutations or the purifying selection against the linked deleterious mutation load [3].

Here we hypothesize that sign variability in fitness effects is necessary for a mutation to exhibit sign inversion. We begin with a brief review of sign-variable mutations in Table 1. We show that they can all be classified into two categories based on whether their effects vary *within* or *between* different lineages. Here, we define a lineage as a complete genealogy of a mutation from its first carrier to its ultimate fixation or loss (Fig. 1, see also [2]). We use computer simulations to demonstrate that both classes of sign variability can produce sign inversion and elucidate the underlying population genetic mechanism for each case. Indeed, we find that all reflect a common mechanistic basis, which helps explain why sign variability is ultimately necessary for sign inversion. We conclude with some thoughts on the breadth of the evolutionary significance of sign inversion and suggestions for future work.

## RESULTS

### Published cases of sign inversion suggest a classification of sign-variable effects

In the studies of Gillespie [5] and Raynes et al. [3] sign variability manifests between individual lineages of the mutation, with some lineages defined by the cost and others by the fitness advantage (Fig. 1C). Gillespie [5] showed that the probability of fixation of a mutation that simultaneously increases the expectation and the variance in offspring number depends on *N*. In fact, when plotted against *N*, Gillespie’s expression for *P*_fix_ exhibits sign inversion (Fig. 1B). Note that higher expectation in offspring number increases the probability that an individual carrier will produce more offspring than non-carriers. Increased variance, however, raises the probability that a carrier leaves no offspring at all. As a result, small lineages of such a mutation are more likely to be lost from the population than same-size non-carrier lineages. Those lineages, that by bad luck fail to reproduce, bear the cost of the mutation. Conversely, lineages with the good luck to survive, despite the early risk, come to enjoy the increased expectation in offspring number associated with the mutation. Similarly, Raynes et al. [3] demonstrated sign inversion in selection on mutations that increase the genome-wide mutation rate (called mutators). When recombination is rare, mutators experience indirect selection via statistical associations with fitness-affecting mutations elsewhere in the genome. Lineages that bear the cost of the mutator mutation become linked with excess deleterious mutations and are readily eliminated from the population by purifying selection. On the other hand, mutator lineages defined by linked beneficial mutations are endowed with the fitness advantage provided by the mutator.

In contrast, in the studies of Nowak et al. [6] and Whitlock et al. [4] sign variability manifests among individual carriers in the same lineage, modulated either by mutation frequency or by time. Nowak et al. investigated the evolution of cooperation in the iterated Prisoner’s dilemma game to show that a strategy called tit-for-tat, or TFT, exhibits sign inversion in competition with a strategy called always defect, or ALLD. Because TFT is the better strategy against other TFT but the worse strategy against ALLD, the fitness effect of a TFT mutation is frequency-dependent. When rare, carriers of the TFT mutation suffer a fitness cost. But TFT lineages that manage to increase in size come to enjoy a frequency-dependent fitness advantage. Meanwhile, Whitlock et al. found sign inversion in selection on “recombiners” - mutations that allow for an increased rate of recombination between individuals [4]. Immediately upon appearance, recombiner carriers suffer recombination load – the fitness cost of breaking up beneficial allele combinations during reproduction. However, over time, recombiner lineages evolve increased robustness to recombination and a correlated, decreased deleterious mutation rate, *U*_*del*_. This latter effect affords surviving recombiner carriers a higher equilibrium mean fitness and, correspondingly, a long-term advantage over the non-recombining individuals [4].

### All classes of sign variability yield sign inversion

To test our hypothesis of the causal role of sign variability and to elucidate the underlying mechanism of sign inversion, we studied the evolution of sign-variable mutations in individual-based simulations. We investigated the fixation probabilities of mutations exhibiting between-lineage variability and within-lineage variability, the latter modulated by either time or frequency. Simulations were conducted over a range of population sizes *N*. Each individual simulation was initiated with a single sign-variable invader and *N*-1 wild-type residents. The invader’s *P*_fix_ was estimated as the fraction of replicate simulations in which it reached fixation.

Our simulations confirm that between-lineage sign variability yields sign inversion (Fig. 2, purple open circles). Based on our understanding of sign variability in the studies of Gillespie [5] and Raynes et al. [3], we simulated an invading mutant that with probability *P*_cost_ initiates a lineage bearing a fitness cost ln(*w*)=1+*s*_cost_ and with probability *P*_ben_ = 1 – *P*_cost_ initiates a lineage endowed with a fitness advantage ln(*w*)=1+*s*_ben_. The resident population has a constant fitness ln(*w*)=1. Note that the concave up shape of the sign-variable invader’s *N*·*P*_fix_ in Fig. 2 matches that observed in the two published examples of between-lineage sign variability (Table 1).

**Fig 2:**
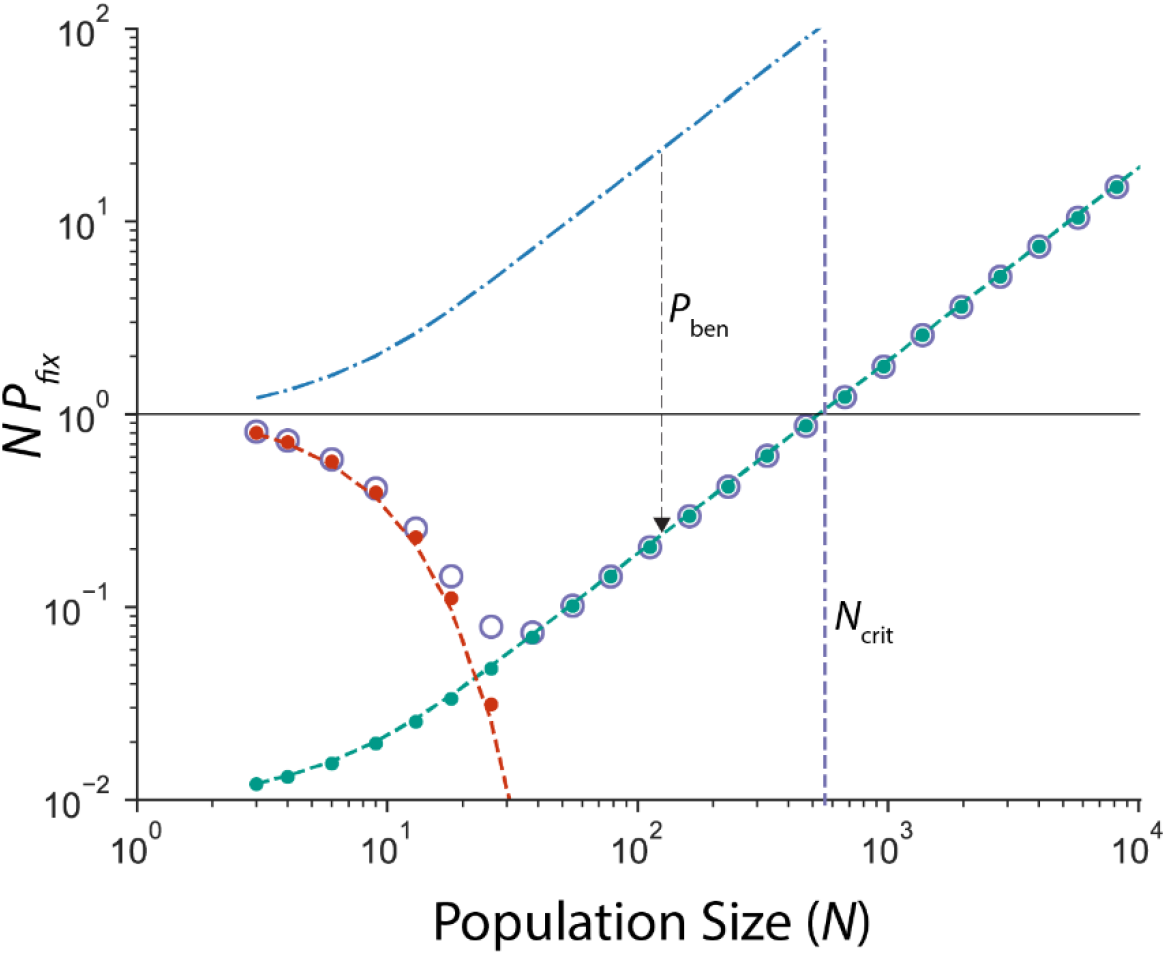
Between-lineage sign variability yields sign inversion. *N*·*P*_fix_ (purple open circles) of the sign-variable invader crosses 1 at *N*_crit_ and can be partitioned into *N*·*P*_fix|cost_ (red, solid circles from simulations, line given by Eq. 2b) and *N*·*P*_fix|benefit_ (teal, solid circles from simulations, line given by Eq. 2c). *N*·*P*_fix_(*N*, *x*=1/*N*, *s*_ben_) is given by the blue dash-dotted line. The downward arrow illustrates that, compared to *N*·*P*_fix_(*N*, *x*=1/*N*, *s*_ben_), *N*·*P*_fix|benefit_ is everywhere reduced by constant *P*_ben_ (see Eq. 2c). Parameter values: *s*_ben_=0.1, *s*_cost_=-0.1, *P*_ben_= 0.01, *P*_cost_ = 0.99. All simulation results are averaged across 10^7^ replicates.

To elucidate the mechanism of sign inversion for between-lineage variability, we followed Raynes et al. [3] by partitioning *N*·*P*_fix_ of the sign-variable mutation into probabilities of fixation in lineages that bear the fitness cost and in lineages endowed with the fitness benefit – hereafter, respectively, *N*·*P*_fix|cost_ and *N*·*P*_fix|benefit_. Since the sign-variable mutation can only fix via one of these two paths,

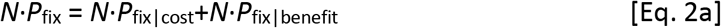

where, again following [3],

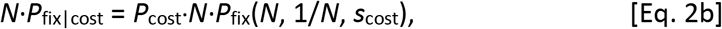

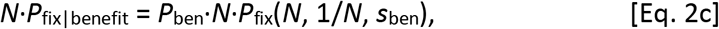

and *P*_fix_(*N*, *x*, *s*) is given by Eq. 1 above. As Fig. 2 illustrates, selection against the cost-bearing lineages pushes *N*·*P*_fix|cost_ below 1 at all *N* > 1. That is, cost-bearing lineages always fare worse than the neutral benchmark. Meanwhile, *N*·*P*_fix|benefit_ increases with *N*, similar to the fixation probability of a lineage-invariant beneficial mutation (blue in Fig. 2). In fact, as Eq. 2c shows *N*·*P*_fix|benefit_ is, simply, the lineage-invariant *N*·*P*_fix_(*N*, 1/*N*, *s*_ben_) reduced everywhere by the probability *P*_ben_ (Fig.2). As a result, while *N*·*P*_fix_(*N*, 1/*N*, *s*_ben_) never crosses 1, *N*·*P*_fix|benefit_ is below 1 at 1 < *N* < *N*_crit_. Sign inversion occurs because *N*·*P*_fix_ is dominated in populations above *N*_crit_ by the higher-than-neutral *N*·*P*_fix|benefit_. But as *N* declines, selection for benefit-endowed lineages is overwhelmed by drift (i.e., *N*·*P*_fix|benefit_ crosses 1) before selection against cost-bearing lineages. As a result, *N*·*P*_fix_ transits below 1 at *N* = *N*_crit_ and is dominated at *N* < *N*_crit_ by the lower-than-neutral *N*·*P*_fix|cost_.

The moderating effect of *P*_ben_ on sign-variable *N*·*P*_fix_ in Eq. 2 illustrates the basic principle of sign inversion, that we will show to be common to all classes of sign variability. In every model of sign variability, mutant fixation probability can only be raised above the neutral benchmark by selection for carriers endowed with the fitness advantage. Yet, because only a fraction of carriers experience the advantage, mutant *N*·*P*_fix_ will always be lower than the corresponding lineage-invariant *N*·*P*_fix_(*N*, *x*, *s*_ben_). Indeed, we will show that in every model of sign variability, mutant *N*·*P*_fix_ is reduced below lineage-invariant *N*·*P*_fix_(*N*, *x*, *s*_ben_) by the probability of *avoiding*, what we will call, the evolutionary hazard of sign-variability (Table 2). We designate this probability 1-*P*_hazard_, where *P*_hazard_ is the probability of succumbing to the hazard. That is, the fixation probability of every sign-variable allele, afforded to it by selection on its benefit-endowed carriers, can be thought of as the product (1-*P*_hazard_)·*P*_fix_(*N*, *x*, *s*_ben_). For example, in the model of between-lineage variability (Fig. 2), the hazard of sign variability lies in producing a deleterious rather than a beneficial lineage, which occurs with probability *P*_cost_. Accordingly, Eq. 2c shows *N*·*P*_fix|benefit_ to be reduced by 1-*P*_hazard_ given by 1-*P*_cost_ = *P*_ben_ (Fig. 2). Note also that lineage-invariant *N*·*P*_fix_(*N*, *x*, *s*_ben_) always necessarily increases with *N* (e.g., Fig.1A). As a result, because 1-*P*_hazard_ (i.e., *P*_ben_) in Fig. 2 is constant in *N, N*·*P*_fix|benefit_ (given by Eq. 2c) has a positive slope at *N*_crit_ and mutant *N*·*P*_fix_ is concave-up. But as we shall see, this population size independence of *P*_hazard_ does not exist in every model of sign-variable fitness.

**Table 2:** Evolutionary hazards of sign variability.

| Table 2: Evolutionary hazards of sign variability |  |  |  |  |  |
| --- | --- | --- | --- | --- | --- |
| Type of Sign Variability | Evolutionary hazard | $1 - P_{\text{hazard}}$ | $1 - P_{\text{hazard}}$ declines faster than $1/N$? | Shape of $N \cdot P_{\text{fix}}$ | Figs. |
| *Between-lineage sign variability* | Failing to produce a beneficial lineage | $P_{\text{ben}}$ | No | Concave-up | 2 |
| *Within-lineage sign variability* |  |  |  |  |  |
| *Positive time-dependence* | Failing to survive until time $t$ while deleterious | $P_{\text{surv}}(\text{until } t)$ | No | Concave-up | 3A, 3B |
| *Negative time-dependence* | Failing to fix before time $t$ while beneficial | $P_{\text{fix}}(\text{before } t)$ | Yes | Concave-down | 3A, S1 |
| *Positive frequency-dependence* | Failing to transit from $1/N$ to $x^*$ while deleterious | $P_{1/N \rightarrow x^*}$ | Yes | Concave-down | 4A, 4B |
| *Negative frequency-dependence* | Failing to transit from $x^*$ to 1 while deleterious | $P_{x^* \rightarrow 1}$ | Yes | Concave-down | 4A, S2 |

We next studied the two models of within-lineage sign variability in which the fitness effect is modulated by either time or frequency (based, respectively, on our understanding of [4] and [6]). In our model of time-dependent sign variability, the sign of the fitness effect conferred on all contemporary carriers within a lineage changes at time *t*. Under positive time-dependent selection, the fitness effect switches from *s*_cost_ to *s*_ben_ (inspired by recombiners in [4] and illustrated in Fig. 1D). Under negative time-dependent selection, the fitness effect switches from *s*_ben_ to *s*_cost_. And similarly, in our model of frequency-dependent sign variability, the sign of the fitness effect conferred on all contemporary carriers within a lineage changes when the mutation crosses threshold frequency *x\**. Again, under positive frequency-dependence (inspired by TFT mutants in [6]), the fitness effect switches from *s*_cost_ below *x\** to *s*_ben_ above *x\**. Under negative frequency-dependent selection, the fitness effect switches from *s*_ben_ to *s*_cost_. In both models, the resident population has a constant fitness ln(*w*)=1.

We find that both models of time-dependent selection exhibit sign inversion (Fig. 3A). *N*·*P*_fix_ of an allele experiencing positive time-dependent selection is concave up (Fig. 3A, purple), similar to that of a recombiner mutation (Ref. [4] and Table 1). In contrast, *N*·*P*_fix_ of an allele experiencing negative time-dependent selection is concave down (Fig. 3A, orange). To understand the mechanism of sign inversion for time-dependent sign variability, we again partitioned *N*·*P*_fix_ into fixation probabilities in carriers that bear the cost, *N*·*P*_fix|cost_, and carriers endowed with the benefit, *N*·*P*_fix|benefit_. Eq. 2a applies again here, as these mutations can fix either while beneficial or while deleterious.

**Figure 3:**
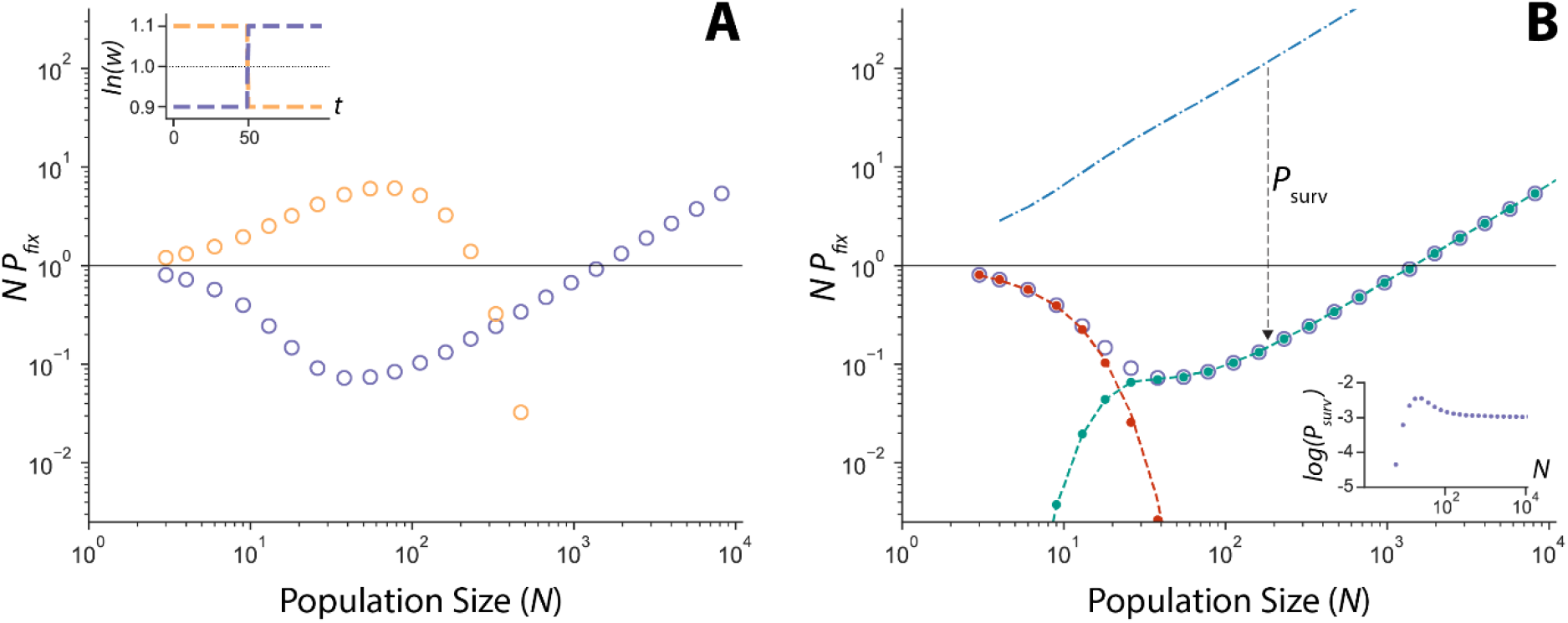
Sign inversion for within-lineage sign variability modulated by time. **A)** Both sign-variable alleles under positive (purple) and negative (orange) time-dependent selection exhibit sign inversion. Simulation results averaged across 10^7^ replicates. **Inset)** Parameter values: *s*_ben_=0.1, *s*_cost_=−0.1, *t*=50. **B)** *N*·*P*_fix_ under positive time-dependent selection (purple open circles, identical to results in panel A) can be partitioned into *N*·*P*_fix|cost_ (red, solid circles from simulations, line given by *N*·*Pfix*(*N*, 1/*N*, *s*_cost_)) and *N*·*P*_fix|benefit_ (teal, solid circles from simulations, line given by Eq. 3). 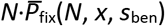 is shown by the blue dash-dotted line (Methods). The downward arrow illustrates that, compared to 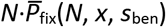, *N*·*P*_fix|benefit_ is everywhere reduced by *P*_surv_(*t=*50) (see Eq. 3). **Inset**) *Psurv*(*t=*50) calculated from simulations in panel A.

Under positive time-dependent selection (purple in Fig. 3A), *N*·*P*_fix|cost_ and *N*·*P*_fix|benefit_ describe, respectively, fixation probabilities before and after time *t*. Reflecting selection against the short-term cost, *N*·*P*_fix|cost_ is below the neutral threshold at all *N* > 1 and is well estimated simply by the lineage-invariant *N*·*P*_fix_(*N*, 1/*N*, *s*_cost_) given by Eq. 1 (Fig. 3B). Meanwhile, *N*·*P*_fix|benefit_ rises with *N* but is again depressed by 1-*P*_hazard_ (Table 2). In this case, because only those lineages that do not succumb to the short-term cost experience the long-term benefit, 1-*P*_hazard_ is given by the probability that a mutant lineage survives until time *t*, or *P*_surv_(*t*). Algebraically, we find

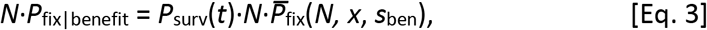

where 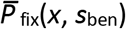 is the average of fixation probabilities given by Eq. 1, weighted by the distribution of mutant frequencies *x* at time *t* in our simulations (see Methods). *N*·*P*_fix|benefit_ given by Eq. 3 crosses from below 1 at *N* < *N*_crit_ to above 1 at *N* > *N*_crit_ (Fig. 3B). Thus, as in the model above, sign inversion occurs because selection for the benefit (represented by *N*·*P*_fix|benefit_) is overwhelmed by drift at a larger *N* than is necessary for drift to neutralize selection against the cost (represented by *N*·*P*_fix|cost_).

Notice also that while 1-*P*_hazard_ in this model – *P*_surv_(*t*) – is inflated by drift at small *N*, it quickly converges to an *N*-independent value (Fig. 3B). Correspondingly, because 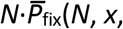 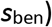 is, as always, increasing with *N* (Fig. 3B), *N*·*P*_fix|benefit_ in Eq. 3 has a positive slope at *N*_crit_ and mutant *N*·*P*_fix_ is concave up. In contrast, in the model of negative time-dependent selection, 1-*P*_hazard_ – now given by the probability that a lineage-invariant beneficial mutant destined to fix does so within the first *t* generations – decreases as *N* increases (Table 2 and Supplementary Information). Moreover, because this 1-*P*_hazard_ declines faster than the reciprocal of *N* (Supplementary Fig. 1), *N*·*P*_fix|benefit_ has a negative slope at *N*_crit_ and, consequently, *N*·*P*_fix_ is concave-down.

Finally, we find that both models of frequency-dependent selection also exhibit sign inversion (Fig. 4). Interestingly, unlike time-dependent selection, both positive and negative frequency-dependent selection yields concave-down *N*·*P*_fix_. Here again we sought to partition *N*·*P*_fix_ into probabilities corresponding to carriers bearing the cost and carriers endowed with the benefit. However, because the fitness effect of every sign-variable mutant switches at some frequency *x\**<1, separating *N*·*P*_fix_ into *N*·*P*_fix|cost_ and *N*·*P*_fix|benefit_ (as in Eq. 2a) is not possible. Put another way, since a mutant lineage must transit past *x\** on the way to fixation, it cannot fix while deleterious under positive frequency-dependence nor while beneficial under negative frequency-dependence. Consequently, we instead partitioned *N*·*P*_fix_ into the product of the probabilities of transiting from frequency 1/*N* to *x\** and the probability of transiting from frequency *x\** to fixation. Algebraically, one can easily show that a neutral mutation will transit from frequency 1/*N* to *x\** with probability 1/(*N*·*x\**), and from *x\** to 1 with probability *x\**. Thus we now write

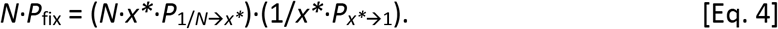

**Figure 4:**
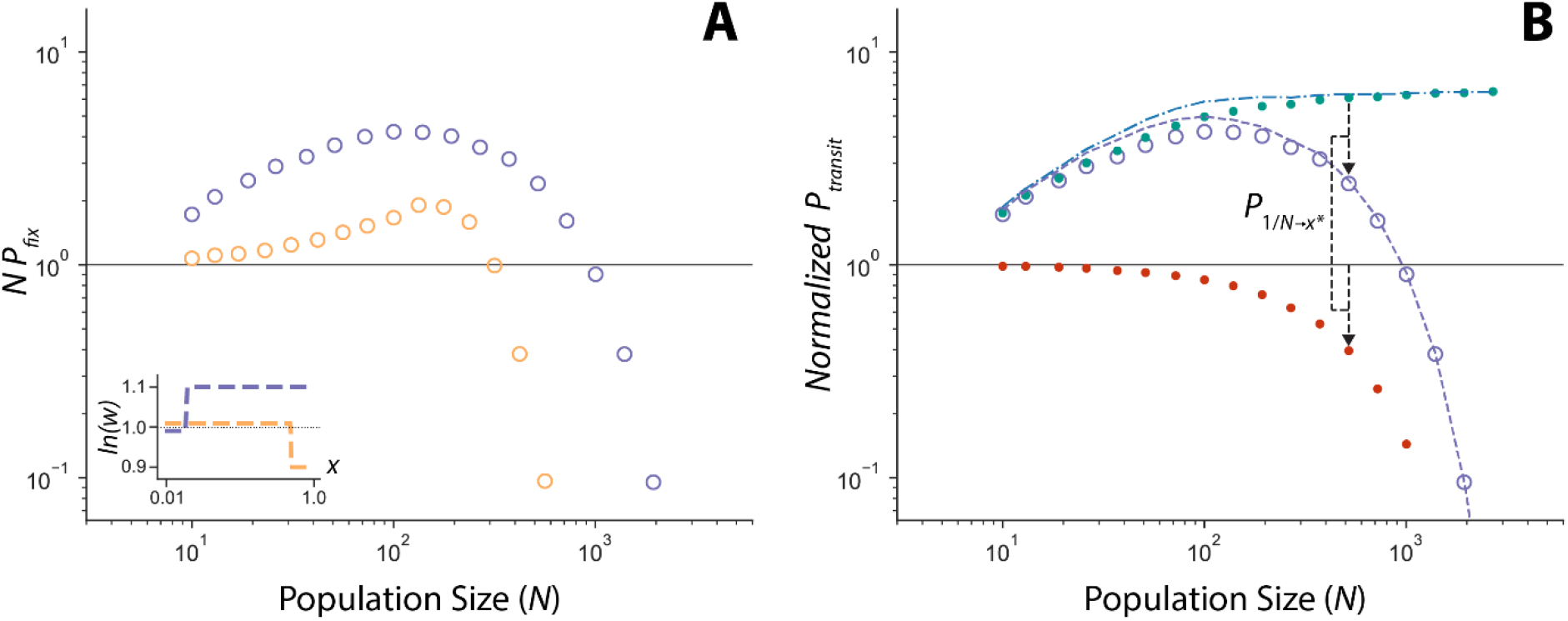
Sign inversion for within-lineage sign variability modulated by frequency. **A)** *N*·*P*_fix_ of sign-variable mutations under both positive (purple circles) and negative (orange circles) frequency-dependent selection exhibit sign inversion. Simulation results are averaged across 10^7^ replicates. **Inset)** Parameter values: for positive frequency-dependence, *s*_cost_ = −0.01, *s*_ben_ = 0.1, *x\** = 0.15; for negative frequency-dependence, *s*_cost_ = −0.1, *s*_ben_ = 0.01, *x\** = 0.85. **B)** The total *N*·*P*_fix_ (purple open circles, identical to results in panel A) of the sign-variable invader under positive frequency-dependent selection can be regarded as the product of *N*·*x\**·*P*_1/*N*→*x\**_ (red solid circles) and 1/*x\**·*P*_*x\**→1_ (teal solid circles). 1/*x\**·*P*_*x\**→1_ is well predicted by 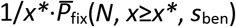 given by the blue dash-dotted line. Purple dashed line is given by the product (*N*·*x\**·*P*_1/*N*→*x\**_)·(1/*x\**·*P*_fix_(*N*, *x*≥*x\**, *s*_ben_)). Downward arrows illustrate that sign-variable fixation probability (purple) is reduced compared to the lineage-invariant fixation probability of a beneficial mutant (blue) by P_1/*N*→x*\**_ (red, here normalized by *N*·*x\**)

As shown next, one of these two transit probabilities can be regarded as 1-*P*_hazard_ and the other as the lineage-invariant *P*_fix_(*N*, *x\**, *s*_ben_), contingent on whether the frequency dependence is positive or negative.

Fig. 4B shows the partitioning of *N*·*P*_fix_ for positive frequency-dependent selection as suggested by Eq. 4. In this this case, the first transit probability in Eq. 4 (*N*·*x\**·*P*_1/*N*→*x\**_) is associated with selection against the mutant’s cost and is below the neutral expectation at all *N*. Indeed, because it describes the probability that a mutant lineage will avoid the hazard of positive frequency-dependent selection (i.e., selective loss while at low frequency) it can be regarded as 1-*P*_hazard_ (Table 2). The second transit probability in Eq.4 (1/*x\**·*P*_*x\**→1_) is associated with selection for the fitness benefit and is consistently above the neutral expectation. Here, as in the models above, sign inversion occurs because selection against the cost and selection for the benefit are overwhelmed by drift at different *N*. In small population in which *N*·*x\**·*P*_1/*N*→*x\**_ is closer to the neutral expectation, *N*·*P*_fix_ is dominated by higher-than-neutral 1/*x\**·*P*_*x\**→1_ and the sign-variable allele behaves most like a beneficial mutation with *N*·*P*_fix_ > 1. However, as *N* increases and selection against the cost overcomes the influence of drift, *N*·*x\**·*P*_1/*N*→*x\**_ begins to decline, which pushes *N*·*P*_fix_ below the neutral expectation.

Considering the two transit probabilities also helps explain the concavity of *N*·*P*_fix_. Because 1/*x\**·*P*_*x\**→1_ (the second factor in Eq. 4) describes the probability of fixation while the mutant is beneficial, it is well approximated by the monotonically increasing lineage-invariant 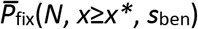 (Fig. 4B). (Note that under the Wright-Fisher construction of our simulations, invading lineages do not necessarily reach 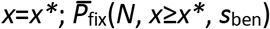 is, thus, the average of fixation probabilities given by Eq. 1,weighted by the distribution of mutant frequencies *x* at which the mutation reaches above *x\**). Meanwhile, 1-*P*_hazard_ given by the first factor in Eq. 4 – i.e., the probability that a mutant reaches *x\** while deleterious – declines with population size. Indeed, similar to the fixation probability of a deleterious mutation (red line in Fig. 1A), it does so faster than 1/*N*. As a result, *N*·*P*_fix_ – the product of a lineage-invariant *P*_fix_ and (1-*P*_hazard_) – declines with population size as well (i.e., is concave down in Fig. 4B).

Sign inversion has an exactly analogous explanation in the case of negative frequency dependent selection (Supplementary information). Here again Eq. 4 applies, but now it is the second factor (1/*x\**·*P*_*x\**→1_) that gives the probability that a mutant lineage escapes the hazard of sign variability, i.e., 1-*P*_hazard_. Meanwhile, the first term *(N*·*x\**·*P*_1/*N*→*x\**_) describes the expected invariant probability of the lineage’s evolutionary success. However as above, 1-*P*_hazard_ in this model drops faster than the reciprocal of *N*, accounting for the concave down shape of *N*·*P*_fix_ seen in Fig. 4A (Supplementary Fig. 2).

## DISCUSSION

We have shown that both within-lineage and between-lineage sign variability in fitness effects can cause sign inversion, thus accounting for all published cases in Table 1. To elucidate the common mechanism of sign inversion, we partitioned *N*·*P*_fix_ of each sign-variable allele into component probabilities associated with fixation of its cost-bearing and benefit-endowed carriers. We found that sign inversion always occurs because selection against the former and selection for the latter exhibit differing sensitivities to drift. In other words, the cost-associated and benefit-associated components of *N*·*P*_fix_ approach the neutral expectation at different *N*. For instance, in our model of positive time-dependence (Fig. 3B) *N*·*P*_fix_ is dominated by selection on benefit-endowed carriers (represented by *N*·*P*_fix_|ben*efit*) at *N*>*N*_crit_. But, selection for benefit-endowed carriers is overwhelmed by drift at a larger *N* than is selection against cost-bearing carriers (represented by *N*·*P*_fix|cost_). That is, *N*·*P*_fix_|ben*efit* approaches 1 at a larger *N* than *N*·*P*_fix|cost_. As a result, *N*·*P*_fix_ transitions below 1 when *N* declines below *N*_crit_. Conversely, *N*·*P*_fix_ of an allele under positive frequency-dependent selection (Fig. 4B) is dominated by selection against its cost-bearing carriers at *N*>*N*_crit_. But as *N* declines, selection against the cost is effectively neutralized by drift before drift can overwhelm selection for the benefit-endowed carriers. Correspondingly, as *N* declines below *N*_crit_, *N*·*P*_fix_ transitions above 1.

The discrepancy between sensitivities to drift while beneficial and deleterious in every case of sign inversion confirms our intuition that sign variability is necessary for sign inversion. After all, sign variability allows carriers of a single mutation to experience both a fitness cost and a fitness benefit, that, in turn, can differentially affect its evolutionary fate at different *N*. In contrast, all carriers of a sign-invariant mutation can only ever experience a fitness cost (if the mutation is deleterious) or a fitness benefit (if the mutation is beneficial). When that singular selective pressure is overpowered by drift, the sign-invariant mutation is rendered neutral in all carriers, precluding sign inversion. Note, though, that while sign variability is necessary, it is not sufficient for sign inversion. Indeed, it can be easily shown that whether the fixation probability of a sign-variable mutation ever crosses the neutral expectation depends on the relative strengths of selection on its cost-bearing and benefit-endowed carriers (see Supplementary Fig. 3 for an example of a between-lineage variable allele that does not exhibit sign inversion). We leave it for future studies to define the conditions under which sign-variable mutations do and do not experience sign inversion.

Our results also help explain the two alternative shapes of *N*·*P*_fix_ observed here and in published cases of sign inversion. Recall that, in every model of sign variability, the mutant’s *N*·*P*_fix_ is reduced below *N*·*P*_fix_(*N*, *x*, *s*_ben_) of a lineage-invariant beneficial mutation by 1-*P*_hazard_. Because the lineage-invariant *N*·*P*_fix_(*N*, *x*, *s*_ben_) monotonically increases with *N* (see Fig. 1A), the shape of sign-variable *N*·*P*_fix_ is determined by the scaling relationship between 1-*P*_hazard_ and *N*. Provided that 1-*P*_hazard_ does not decline faster than the reciprocal of population size sign-variable *N*·*P*_fix_ must eventually rise above the neutral threshold with *N*, resulting in a concave up *N*·*P*_fix_ (Table 2). Conversely, in cases in which 1-*P*_hazard_ does decline faster than 1/*N*, sign-variable *N*·*P*_fix_ must eventually transition below the neutral threshold with increasing *N*, resulting in a concave down *N*·*P*_fix_ (Table 2).

Most generally, our results demonstrating the causal role of sign variability in sign inversion significantly expand our understanding of the latter’s evolutionary importance. Sign variability in nature is surely not limited to the four published cases of sign inversion.

Previously, Graves and Weinreich [2] showed that variation in fitness among allele’s carriers could be facilitated by their interactions with environmental, social, or genetic factors. For example, time-dependent within-lineage sign variability could be modulated by temporal changes in the lineage’s environment. Alternatively, interactions with other individuals in a population could give rise to frequency-dependent (as in [6]) or density-dependent within-lineage sign variability. Meanwhile, between-lineage sign variability may emerge in any scenario in which the fitness effects of a mutation depend on genetic variation at other loci. For example, mutations in some modifier genes are known to experience indirect selection through associations with beneficial and deleterious mutations elsewhere in the genome. Modifiers control a diversity of organismal features [7] such as ploidy, dominance, pleiotropy, migration rate, and mating preference, all of which could be potentially susceptible to sign inversion. Indeed, we have now seen sign inversion in models of ploidy (Raynes and Weinreich, in prep) and of bet-hedging (Weissman et al., in prep). We strongly suspect that future studies will discover sign inversion in a variety of biological models revealing a much more general influence of population size on the direction of natural selection.

## METHODS

### Simulations

We model asexual populations of constant size, *N*, evolving in discrete, non-overlapping generations under the influence of selection and drift as the only evolutionary forces (i.e., without mutation). Each simulation starts with two lineages: one comprising a single sign-variable invader and the other comprising *N*-1 sign-invariant residents. Simulation ends when the sign-variable lineage is either fixed or lost from the population.

Every generation, the population reproduces according to the Wright-Fisher model [8]. The size of each lineage in the next generation is randomly drawn from a binomial distribution with expectation *N*·*x*·*w*, where *x* is the frequency of the lineage in the previous generation and *w* is the relative fitness of the lineage (i.e., lineage’s fitness divided by the average fitness of the population). The fitness of the invariant residents is always ln(*w*)=1.0. The fitness of the sign-variable invaders is determined as follows.

#### Between-lineage sign variability

At the beginning of each replicate simulation, the fitness of the sign-variable invader is chosen randomly to be either ln(*w*) = 1+*s*_cost_ or ln(*w*)=1+s_ben_ with probabilities *P*_cost_ and *P*_benefit_ respectively. The invader’s fitness remains unchanged in all its descendants until the end of the simulation.

#### Within-lineage sign variability

At the beginning of each replicate simulation, the fitness of the sign-variable invader is set to either ln(w) = 1+*s*_cost_ for mutations under negative time-and frequency-dependent selection or ln(*w*) = 1+*s*_ben_ for mutations under positive time-and frequency-dependent selection. It remains unchanged in all its descendants until either time *t* (under time-dependent selection) or until its frequency exceeds critical frequency *x\** (under frequency-dependent selection). The fitness of all contemporary sign-variable carriers is then instantaneously changed from ln(*w*) = 1+*s*_ben_ to ln(*w*) = 1+*s*_cost_ for models of negative time- and frequency-dependent selection or vice versa for models of positive time- and frequency-dependent selection.

To calculate the expected lineage-invariant fixation probabilities 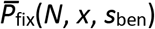 (for positive *time*-dependent selection) and 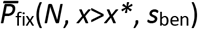 (for positive *frequency*-dependent selection) we recorded the invader frequency *x* in every simulation that reached either time *t* or frequency *x\**. In the former case, we assayed the invader frequency in generation *t*+1. In the latter, we assayed invader frequency in the generation immediately after it surpassed *x\**. (Note that because of the Wright-Fisher construction of our simulations, a lineage wouldn’t necessarily reach exactly frequency *x\**.) We then calculated 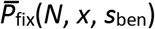 as the average *P*_fix_(*N*, *x*, *s*_ben_) given by Eq. 1 for every *x* in the distribution.

Note that our estimate of 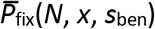 for positive frequency-dependence in Fig. 4 does not take into account that a frequency-dependent mutation has an increased probability of extinction from frequency *x\**, i.e., *P*_*x\**→0_, compared to a lineage-invariant mutation. As a result, it can slightly overestimate the beneficial transit probability *P*_*x\**→1_, especially at small *N* at which the probability of drifting below *x\** is at its highest.

The frequency distribution, *x*, of lineages that successfully reached *x\** was also used to normalize transit probabilities *P*_1/*N*→*x\**_ and *P*_*x\**→1_ in models of frequency-dependent selection. Given that the expected fixation probability of a neutral mutation equals its starting frequency, the neutral expectation for *P*_*x\**→1 = *x\** (therefore, in Eq. 4 the second factor 1/*x\**·*Px\**→1_ contains the reciprocal 1/*x\**). For our data, we calculated this expectation as the average frequency 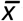 for all lineages that reached x*. If the expected neutral *P*_*x\**→1_ = *x\** and the expected neutral *P*_1/*N*→1_ = 1/*N*, it follows that the neutral expectation for *P*_1/*N*→*x\**_ = (1/*N*)/ *P*_*x\**→1_ = (1/*N*)/*x\** (therefore, in Eq. 4 the first factor *N*·*x\**·*P*_1/*N*→*x\**_ contains the reciprocal *N*·*x\**). For our data, we calculated the neutral expectation for *P*_1/*N*→*x\**_ as 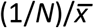.

Simulation code was written in Julia 1.0 and is available at https://github.com/yraynes.

## Acknowledgements

Simulations were performed on the computing cluster of the Computer Science Department at Brown University. The work was supported by the National Science Foundation grants DEB-1556300 and 1736253 to D.M.W. and DEB-2014943 to C.L.B.

## References

1. Kimura, M., On The Probability Of Fixation Of Mutant Genes In A Population. Genetics, 1962. 47: p. 713–719.

2. Graves, C.J. and D.M. Weinreich, Variability in fitness effects can preclude selection of the fittest. Annu Rev Ecol Evol Syst, 2017. 48(1): p. 399–417.

3. Raynes, Y., et al., Sign of selection on mutation rate modifiers depends on population size. Proc Natl Acad Sci U S A, 2018. 115(13): p. 3422–3427.

4. Whitlock, A.O., et al., An Evolving Genetic Architecture Interacts with Hill-Robertson Interference to Determine the Benefit of Sex. Genetics, 2016. 203(2): p. 923–36.

5. Gillespie, J.H., Natural Selection for within-Generation Variance in Offspring Number. Genetics, 1974. 76(3): p. 601–606.

6. Nowak, M.A., et al., Emergence of cooperation and evolutionary stability in finite populations. Nature, 2004. 428(6983): p. 646–50.

7. Otto, S.P., Evolution of Modifier Genes and Biological Systems, in The Princeton Guide to Evolution. 2013: 2013.

8. Ewens, W., Mathematical Population Genetics. 2004, New York: Springer.

