## Supplementary Information for "When good mutations go bad: how population size can change the direction of natural selection"

### 1. Mechanics of sign inversion under negative time-dependent selection.

$N \cdot P_{fix}$ of the mutation with a short-term benefit and a long-term cost transitions from above 1 at $N < N_{crit}$ to below 1 at $N > N_{crit}$ (Supplemental Fig. 1). Its concave-down shape matches that observed for non-recombiners in Whitlock et al. [1]. Accordingly, non-recombiners in [1] are under negative time-dependent selection – they experience a short-term advantage relative to the resident recombiners but eventually develop a relative fitness cost, once recombiners evolve a lower $U_{del}$. To investigate the mechanism of sign inversion negative time-dependent selection, we again partitioned the total $N \cdot P_{fix}$ of the sign-variable mutation into a sum of $N \cdot P_{fix|cost}$ and $N \cdot P_{fix|benefit}$, representing probabilities of fixation while deleterious and beneficial respectively.

$N \cdot P_{fix|benefit}$ describes the fixation probability while beneficial. Given that every sign-variable lineage is beneficial at the outset, we write $N \cdot P_{fix|benefit} = P_{fix}(t < 50) \cdot N \cdot P_{fix}(N, 1/N, s_{ben})$. Here, $P_{fix}(t < 50)$ represents the probability that a lineage-invariant beneficial mutation destined to fix, does so in the first $t$ generations after appearance (i.e., $P_{fix}(N, 1/N, s_{ben})$ in the first $t$ generations divided by the total $P_{fix}(N, 1/N, s_{ben})$).

The inset of Supplemental Fig. 1 shows $P_{fix}(t < 50)$ calculated in simulations of a lineage-invariant invader with a selective advantage $s_{ben}$. In this case, because those lineages that fail to fix before $t$ become deleterious, $P_{fix}(t < 50)$ can be regarded as 1-$P_{hazard}$. Critically, because the time to fixation of a beneficial mutation increases with $N$ [2, 3], $P_{fix}(t < 50)$ declines with $N$ and indeed, it does so faster than 1/$N$ (Supplementary Fig. 1 Inset). As a result, in small populations in which fixation is relatively quick $N \cdot P_{fix|benefit}$ effectively mirrors $P_{fix}(N, 1/N, s_{ben})$. However, as

$N$ increases, $N{\cdot}P_{fix|benefit}$ begins decreasing with $P_{fix}(t < 50)$ and eventually drops below the neutral expectation (Supplementary Fig. 1).

Meanwhile, $N{\cdot}P_{fix|cost}$ corresponds to the period after $t$, when the fitness of sign-variable carriers drops below that of the resident (Supplementary Fig. 1). Note, though, that in this case, sign-variable lineages that survive but do not fix in the first $t$ generations, are likely to still be enriched by selection before experiencing the fitness cost. Thus, the $N{\cdot}P_{fix|cost}$ reflects the probability that the sign-variable mutation, having previously risen to some frequency *x,* can drift to fixation even after it has become deleterious. Accordingly, it can be well estimated by Eq. 3 from the main text: $N{\cdot}P_{fix|cost} = P_{surv}(t){\cdot}N{\cdot}\overline{P}_{fix}(N, x, s_{cost})$, where $P_{surv}(t)$ is the probability of surviving but not fixing in the first $t$ generations while beneficial (assessed in simulations). $N{\cdot}P_{fix|cost}$ is below 1 at most $N$ but does exceed 1 at some intermediate $N$ (Supplementary Fig. 1). Although, as selection becomes more efficient with $N$ and the probability of drifting to fixation declines, $N{\cdot}P_{fix|cost}$ eventually declines as well, crossing below 1 at $N_{crit}$.

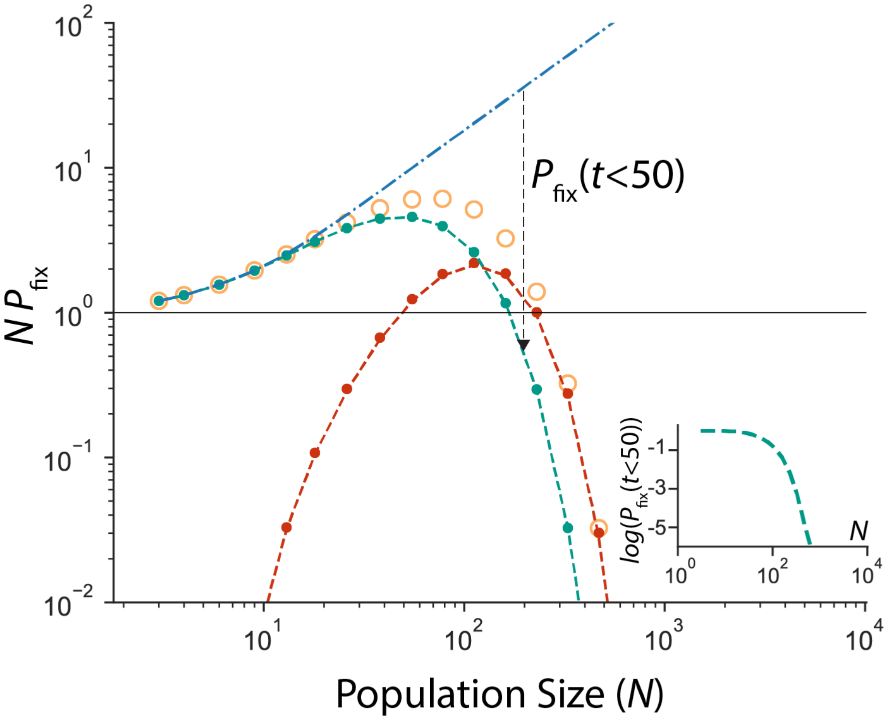


**Supplemental Fig. 1: Sign inversion under negative time-dependent selection.** $N{\cdot}P_{fix}$ (orange open circles) can be partitioned into $N{\cdot}P_{fix|benefit}$ (teal, solid circles from simulations, line given by $P_{fix}(t < 50){\cdot}N{\cdot}P_{fix}(N, 1/N, s_{ben})$) and $N{\cdot}P_{fix|cost}$ (red, solid circles from simulations, line given by $P_{surv}(t){\cdot}N{\cdot}\overline{P}_{fix}(x,$

$s_{cost}$)). $N \cdot P_{fix}(N, x=1/N, s_{ben})$ of the lineage-invariant beneficial mutation is shown as a blue dash-dotted line. The downward arrow illustrates that, compared to $N \cdot \overline{P}_{fix}(N, 1/N, s_{ben})$, $N \cdot P_{fix|benefit}$ is everywhere reduced by $P_{fix}(t<50)$. Parameter values: $s_{ben}$=0.1, $s_{cost}$=-0.1, $t$=50. All simulation results are averaged across $10^7$ replicate simulations. **Inset**: $P_{fix}(t < 50)$ calculated in simulations conducted as described in Methods except with a lineage-invariant invader with fitness $\ln(w)=1+s_{ben}$).

**2. Mechanics of sign inversion under negative frequency-dependent selection.**

To elucidate the mechanism of sign inversion for negative frequency-dependent selection we again partitioned $N \cdot P_{fix}$ into the product of the two transit probabilities ($N \cdot x^* \cdot P_{1/N \rightarrow x^*}$) and ($1/x^* \cdot P_{x^* \rightarrow 1}$). As Supplemental Fig. 2 shows, the probability of transit from $1/N$ to $x^*$ ($N \cdot x^* \cdot P_{1/N \rightarrow x^*}$) is higher than the corresponding probability for a neutral mutation at all $N >$ 1, reflecting the fitness advantage enjoyed by mutant carriers below $x^*$. On the other hand, above $x^*$, the sign-variable mutation becomes deleterious, and must drift to fixation against the direction of natural selection. Correspondingly, the probability of transit from $x^*$ to 1 ($1/x^* \cdot P_{x^* \rightarrow 1}$) is below the neutral expectation at most $N$ and only approaches it in small populations. As a result, in populations below $N_{crit}$, $N \cdot P_{fix}$ is above 1, dominated by the better-than-neutral $N \cdot \mathrm{x}^* \cdot P_{1/N \rightarrow f}$ , but drops below 1 as $N$ increases above $N_{crit}$.

Thus, the concavity of $N \cdot P_{fix}$ here is again explained by the relationship between $N$ and 1-$P_{hazard}$. Supplemental Fig. 2 confirms that the transit probability $N \cdot x^* \cdot P_{1/N \rightarrow x^*}$ effectively mirrors the fixation probability of a lineage-invariant beneficial mutant $N \cdot P_{fix}(N, 1/N, s_{ben})$. This is expected, given that for a beneficial mutation the probability of reaching any frequency high enough to escape drift equals its fixation probability (the discrepancy between the two is due to $P_{1/N \rightarrow x^*}$ being normalized by $1/(N \cdot x^*)$ and $P_{fix}(N, 1/N, s_{ben})$ being normalized by $1/N$). Meanwhile, the hazard of negative frequency-dependent selection is failing to fix even after reaching frequency $x^*$. Thus, the transit probability from $x^*$ to 1, $1/x^* \cdot P_{x^* \rightarrow 1}$, can be regarded as 1-$P_{hazard}$ in this model. Supplementary Fig. 2 shows that this probability, similarly to 1-$P_{hazard}$ under positive frequency-dependent selection, declines faster than $1/N$. Correspondingly, $N \cdot P_{fix}$ – effectively the product of monotonically (but slowly) increasing $N \cdot P_{fix}(N, 1/N, s_{ben})$ and monotonically (but rapidly) declining 1-$P_{hazard}$ – is concave down.

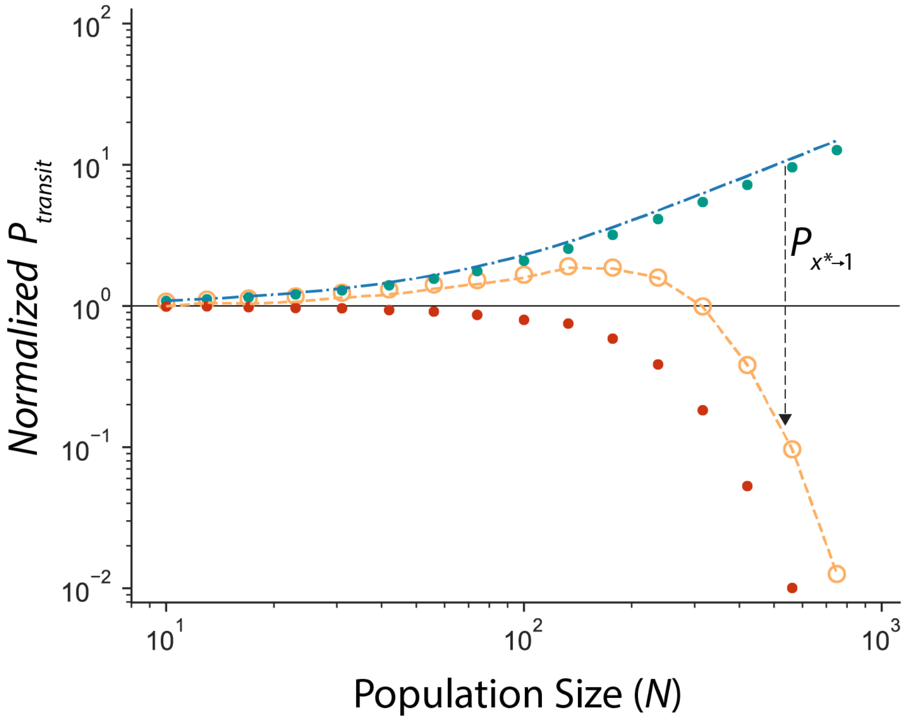


**Supplemental Fig. 2: Sign inversion under negative frequency-dependent selection.** $N \cdot P_{\text{fix}}$ (orange open circles) of the sign-variable invader can be represented as the product of two transit probabilities: $N \cdot x^* \cdot P_{1/N \to x^*}$ and $1/x^* \cdot P_{x^* \to 1}$ (teal and red solid circles respectively). $N \cdot P_{\text{fix}}(N, 1/N, s_{\text{ben}})$ of a lineage-invariant beneficial mutation is shown with the blue dot-dashed line. The product $N \cdot P_{\text{fix}}(N, 1/N, s_{\text{ben}}) \cdot P_{x^* \to 1}$ is given by the dashed orange line. Downward arrow illustrates that sign-variable fixation probability (orange) is reduced compared to the lineage-invariant fixation probability (blue) by $P_{x^* \to 1}$ (red). Parameter values: $s_{\text{cost}}$ = -0.1, $s_{\text{ben}}$ = 0.01, $x^*$ = 0.85. All simulation results are averaged across $10^7$ replicate simulations.

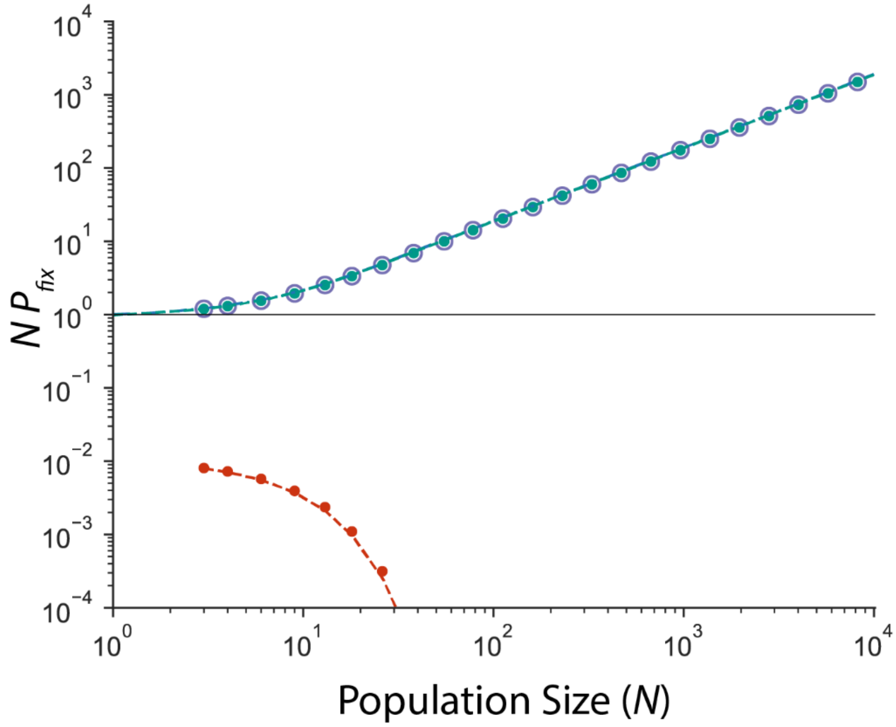


**Supplemental Fig. 3: Sign variability is not sufficient for sign inversion.** Between-lineage sign variability may not yield sign inversion when $P_{ben} > P_{cost}$. The sign-variable $N{\cdot}P_{fix}$ (purple circles) never crosses the neutral expectation (horizontal line). As in the main text, $N{\cdot}P_{fix}$ is partitioned into $N{\cdot}P_{fix|cost}$ (red, solid circles from simulations, line given by Eq. 2b in the main text) and $N{\cdot}P_{fix|benefit}$ (teal, solid circles from simulations, line given by Eq. 2c in the main text). Note that $N{\cdot}P_{fix|benefit}$ never crosses 1 except at $N$=1, at which point the sum of $N{\cdot}P_{fix|cost}$ and $N{\cdot}P_{fix|benefit}$ is exactly 1. Parameter values: $s_{ben}$=0.1, $s_{cost}$=-0.1, $P_{ben}$= 0.99, $P_{cost}$ = 0.01. All simulation results are averaged across $10^7$ replicates.

1. A. O. Whitlock, K. M. Peck, R. B. Azevedo, C. L. Burch, An Evolving Genetic Architecture Interacts with Hill-Robertson Interference to Determine the Benefit of Sex. *Genetics* **203**, 923-936 (2016).
2. M. M. Desai, D. S. Fisher, Beneficial mutation selection balance and the effect of linkage on positive selection. *Genetics* **176**, 1759-1798 (2007).
3. H. Uecker, J. Hermisson, On the fixation process of a beneficial mutation in a variable environment. *Genetics* **188**, 915-930 (2011).
